# Biochemical Indicators of Atlantification and Diapause Strategy in Arctic Copepods Point to a Decrease in Copepod-mediated Carbon Sequestration

**DOI:** 10.64898/2026.07.13.738257

**Authors:** Jiwoon Hwang, Mathieu Lutier, Khuong V Dinh, Katrine Borgå, Bethanie Edwards

## Abstract

Arctic ecosystems are critically endangered by rising temperatures and changing hydrography, especially the intrusion of increasingly warm water from the Atlantic Ocean known as Atlantification. In addition to housing fragile biodiversity, Arctic copepods and their lipids play a crucial role in cycling carbon by transporting carbon into the deep ocean through their diapausing behaviors. Here, we explored the lipidomes of the Arctic copepod *Calanus glacialis*, collected from three fjords around Svalbard during November 2022 when *C. glacialis* are known to be in diapause. These three field sites provide a natural laboratory experiment, as they are influenced by different water masses with varying degrees of Atlantic water, and experience vast differences in sea ice coverage over the year. These environmental differences were clearly reflected in the lipidomic analysis, with stations influenced most by Atlantic Warm Water having the lowest total lipid concentrations and the lowest accumulation of storage lipids necessary for entering diapause. Membrane lipids were a significant proportion of the Svalbard copepod lipidomes, with the highest ratios observed at the Atlantified site. The high membrane lipid and high triacylglycerol concentrations were interpreted as signs of active feeding. This was further corroborated by fatty acid composition analysis, which revealed dietary biomarkers of carnivory at Atlantified sites. The copepods from the site most insulated from Atlantic influence had more than double the amount of storage lipids per individual and fatty acids associated with diatom biomass, indicating assimilation in the spring. Ultimately, the decrease in lipid content observed in association with Atlantification around Svalbard will impact diapause patterns, as *Calanus* species need 20-30% more WE to successfully complete diapause. In turn, this will impact the magnitude of carbon sequestration through the seasonal lipid pump, not to mention having radiating effects through the Arctic food web where *Calanus glacialis* plays an important role.

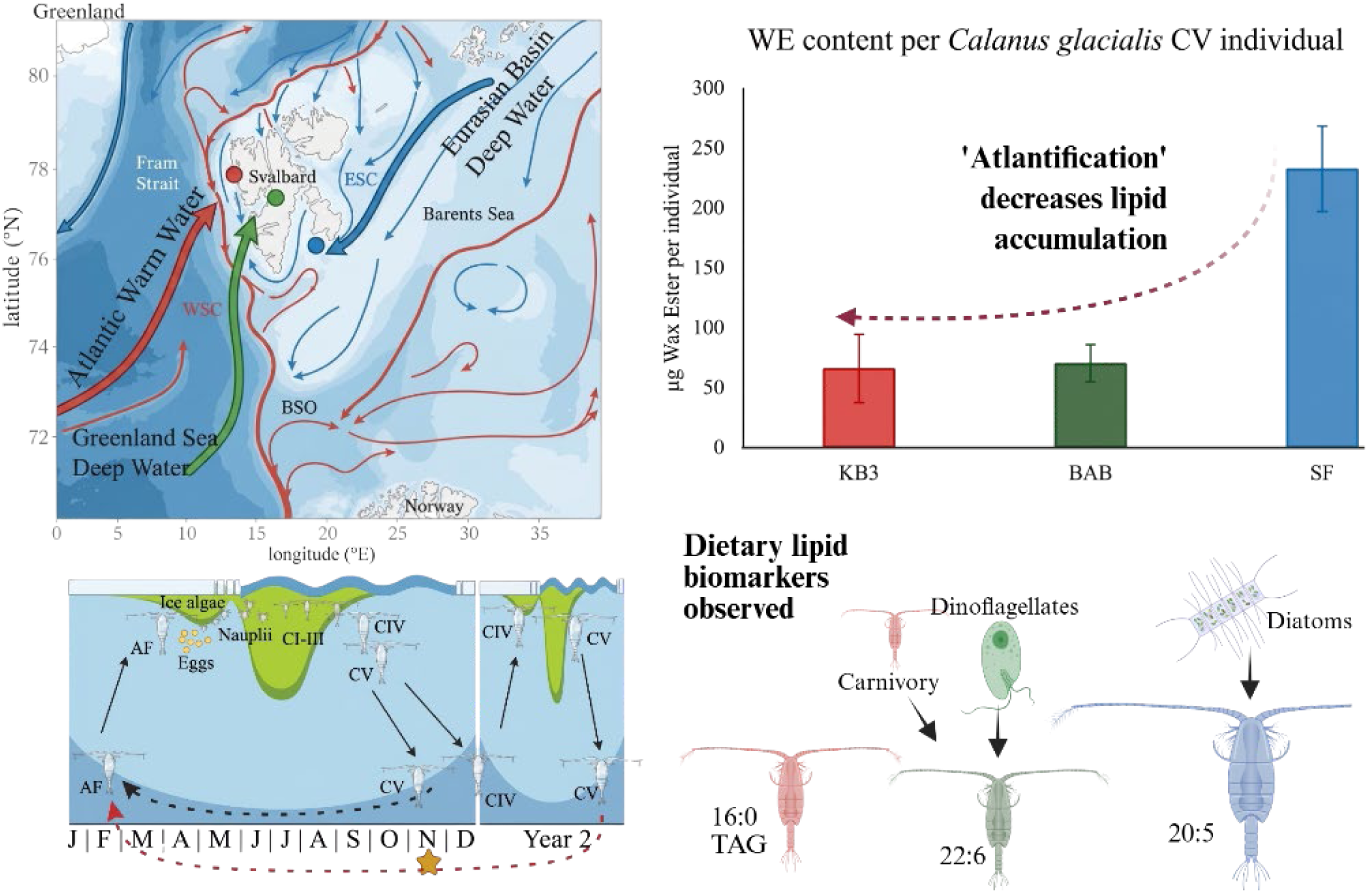

## Introduction

The Arctic is well-recognized as a region of immense environmental significance and a harbinger for climate change. It is one of the most rapidly and intensely changing areas on Earth, with warming temperatures and declining sea ice having cascading effects on the ecosystem [1–6]. A key driver of these changes is the process of “Atlantification,” encompassing both the northward intrusion of warmer, saltier Atlantic waters and *in situ* warming and sea ice decline, which has already reshaped Arctic hydrography, sea ice dynamics, biological communities, and bloom phenology, and is projected to further alter ecosystem structure and function in coming decades [7–11]. Copepods play a critical intermediary role in these polar marine ecosystems, where they contribute significantly to carbon cycling through diel vertical migration and ecological processes by acting as key links between primary producers and higher trophic levels, including birds, mammals, and vital fisheries that depend on these microorganisms (Record et al., 2018 and the references therein) [12].

Beyond their trophic roles, polar copepods contribute considerably to deep sea carbon sequestration through the seasonal lipid pump (SLP) [13–17]. The SLP relies on copepods that have accumulated lipids during the spring bloom transporting carbon to deeper waters (below the permanent pycnocline) during their winter dormancy in a process called diapause. Zooplankton also passively transport carbon to depth through the vertical flux of fecal pellets which in the North Atlantic is estimated to be as large as the sinking flux of particulate organic carbon at similar depths [15]. Wax esters, the dominant lipid class in diapausing copepods, serve as essential long-term energy reserves [18–22]. These lipid compounds have been described as “batteries” charged during productive spring and summer months, sustaining copepods through prolonged periods of overwintering and starvation [12]. Wax esters (WE) are chemically simple compounds that only contain carbon, hydrogen, and oxygen. As such, carbon export via the seasonal lipid pump does not follow the traditional Redfield ratio; i.e., does not deplete the surface ocean of nitrogen and phosphorus. Additionally, these lipid-rich reserves enrich the Arctic food web with energy-dense organic material during seasons of low primary production.

Diapause is a flexible life history strategy for overwintering; its timing, duration, and even occurrence can vary depending on environmental conditions [12, 23–32]. While studies have explored the lipid accumulation window [32, 33], predictive modeling of copepod diapause patterns [12, 23, 24, 30, 34–37], transcriptomic analyses [38–44], and enzyme activity [45], the molecular controls on diapause are still largely unresolved. Approximately 11% of Arctic seas exhibit earlier phytoplankton blooms, predominantly in the lower latitudes, with the annual maximum advancing by as much as 30 – 50 days [4, 46]. Such shifts in bloom timing threaten the “synchrony of phenology”, or the delicate alignment between phytoplankton productivity and copepod feeding. Given their correlation to physiological regulation mechanisms such as buoyancy [47, 48], reproductive maturation [49], and adaptations to cold, high-pressure conditions in deep waters [48], lipids have been proposed as key biochemical signals in diapause, potentially influencing diapause initiation and duration [19]. Lipids also serve as dietary biomarkers, shedding light on feeding ecology and trophic interactions [21, 50–53]. Thus clarifying biochemical signals of diapause is essential for accurately predicting how climate-driven changes may impact carbon fluxes and the distribution of lipid-rich ecosystems at high latitudes [54].

Historically, Arctic copepod lipid research has focused on the cosmopolitan *Calanus finmarchicus* [24, 38, 55–59], leaving significant gaps in our understanding of *Calanus glacialis*, which constitutes ∼80% of Arctic shelf zooplankton biomass [49]. *C. glacialis* is particularly vulnerable to ocean warming due to its relatively shallow diapausing depth [19] and reliance on sea ice algae as a food source during early spring [60, 61]. This vulnerability is further compounded by Atlantification, which is driving the northward expansion of *C. finmarchicus*, a sub-Arctic congener predicted by models to displace *C. glacialis* in warming Arctic waters. Although the capacity of *C. finmarchicus* to sustain its population in the Arctic is still an open question [7, 9, 11, 23], as are the sensitivity and adaptability of *C. glacialis* to increased Atlantic influence. A decline in *C. glacialis* populations is hypothesized to have far-reaching consequences for Arctic biodiversity and ecosystem stability, given its dominant role in the region [34, 35]. For instance, *C. finmarchicus* is smaller-bodied and contains substantially lower lipid reserves than *C. glacialis* [19, 62]. A shift toward *C. finmarchicus*-dominated communities could therefore reduce energy transfer to higher trophic levels, diminish the magnitude of carbon sequestered via the seasonal lipid pump, and put undue stress on populations of the little auk (*Alle alle*), a seabird which feeds exclusively on *C. glacialis* and flies long distances to do so, even when *C. finmarchicus* is readily available [63, 64]. Consequently, a deeper understanding of the life history strategies and environmental sensitivity of *C. glacialis* is essential for predicting broader ecological implications. Here, we investigate the meta-lipidome of *C. glacialis* from three distinct Arctic fjords around Svalbard during wintertime: Kongsfjord (KB3), characterized by minimal to no sea ice; Storfjorden (SF), with extensive ice cover; and Billefjord (BAB), representing intermediate ice conditions. We hypothesize that changes in hydrographic and thermal conditions will be reflected in the lipidomic composition of wintertime copepods. To achieve comprehensive lipid characterization, we utilized untargeted Orbitrap-based lipidomics to analyze the intact lipid pool in detail. Whereas many previous copepod lipid studies have employed fatty acid methyl ester (FAME) analyses or lipid sac volume estimates (*i.e.*, prosome length), providing valuable insights into bulk lipid composition, our approach advances these by enabling a more detailed structural characterization of intact lipid species. Moreover, by comparing lipidomes across fjords and life stages, we explore how lipid profiles vary with environmental and physiological influences, contributing to a deeper understanding of copepod adaptive life strategies in rapidly changing Arctic ecosystems. Lastly, we provide back-of-the-envelope estimates of how severe Atlantification scenarios could diminish the contribution of *C. glacialis* to the seasonal lipid pump, framing these biochemical observations within their broader implications for Arctic carbon cycling.

## Methods

### Experimental Design

Copepods were collected over a 5-day period (November 13–17, 2022) aboard the R/V *G.O. Sars* as part of the Nansen Legacy Project. Samples were obtained using a 180 µm mesh multinet, deployed from the fjord bottom to 100 m above the seafloor. Individuals were morphologically identified to stage and species under a dissecting microscope. Developmental stages were the copepodite stages IV and V of *Calanus glacialis*. Species-level identification relied on prosome segmentation and morphology. *Calanus hyperboreus* was readily distinguished from other *Calanus* species by its larger body size and the presence of a spine at the tip of the prosome. *C. glacialis* and *C. finmarchicus* were tentatively distinguished by antennal coloration, with *C. glacialis* typically exhibiting red antennae and *C. finmarchicus* paler antennae, a method considered reliable in Svalbard fjords [64, 65], though it can overlap between the two species. Genetic confirmation was not available for this sample set, and species assignment should therefore be considered provisional. Individuals of *C. hyperboreus* were also identified but were excluded from this study because of their sparse abundance across our dataset.

Lipid samples of the surface microbial community were collected from 10- and 50-meter depths at all three fjords. Samples were collected by filtering 1 L of seawater through a 0.7 μm GF/F filter. Filters were flash frozen in liquid nitrogen and transported on dry ice.

### Lipid Extraction and Analysis

For each fjord and developmental stage, nine copepods were analyzed. Individuals were extracted in groups of three, yielding three pooled replicates per fjord-stage pair. (Supplementary Table S1). Each replicate was homogenized in a Bligh and Dyer mixture using glass beads to disrupt exoskeleton and cellular membranes, improving lipid recovery, following methods outlined in Hwang and Edwards, 2025 [66]. Each extraction had 10 µL of EquiSPLASH LIPIDOMIX® Quantitative Mass Spec Internal Standard (Avanti Polar Lipids, Alabaster, AL, USA) added as an internal standard.

UPLC separation and mass spectrometric analysis also followed the methods described in Hwang and Edwards, 2025 [66]. Peak identification and annotation were performed using LOBSTAHS (Lipid and Oxylipin Biomarker Screening Through Adduct Hierarchy Sequences) and MS-DIAL. LOBSTAHS relies on adduct hierarchy and retention time, while MS-DIAL uses MS/MS fragment libraries. Specifically, the Svalbard copepod meta-lipidome was queried against the MS-DIAL V17 positive database and our *in silico* WE database [66]. While annotation outputs differed due to database structure, peak area integration was comparable across platforms. Where annotations conflicted, manual inspection of the MS2 fragmentation and adduct hierarchy was used to determine the final identity. The data reduction table is presented as Supplementary Table S2.

### Absolute storage lipid quantification

LC-MS peak areas of WE and TAGs were converted to molar quantities using corresponding 5-point WE and TAG standard curves fitted through the origin, followed by conversion to carbon mass based on elemental composition. Carbon mass per compound was calculated by multiplying molar quantity by the number of carbon atoms and the atomic mass of carbon (12.011 g mol⁻¹). Compound-level carbon masses were summed across relevant lipid species within each sample to obtain *∑TAG, ∑WE,* and *∑WE+TAG*.

Our assumptions included stable population size, unchanged diapause depth and duration, and baseline Arctic overwintering biomass from values derived from Tande and Henderson (1988) and Jónasdóttir et al. (2015) [15, 67]. Values of contribution to the biological carbon pump were calculated by applying a proportional scaling factor to Eastern Norwegian Sea estimates (chosen for their geographical proximity to our sample site) from Jónasdóttir et al. (2015) [15]. The scaling factor was defined as the ratio of carbon measured in the present study to the corresponding carbon values reported in that study. Contribution to the biological carbon pump was assumed to be the total respired carbon of the copepods at depth. For consistent comparison with our reference values and between biological stages, our calculations were based on carbon derived from wax ester in CV individuals only. Amounts of carbon in ktonC per year were computed by multiplying the carbon flux over the given area of the Eastern Norwegian Sea [15]: 1.82E+11 m^2^.

### Environmental Data

Water column data for our sample sites were collected using a CTD deployed at the same time and depth as the multinet. Salinity and temperature were measured using a Sea-Bird Electronics CTD system, dissolved oxygen with an SBE 43 sensor, and fluorescence with a Chelsea Aqua 3 chlorophyll fluorometer. Pressure is used as a proxy for depth (dbar ≈ m).

Chlorophyll-a satellite data was derived from the ESA Ocean Colour Climate Change Initiative (OC-CCI), sea ice concentration from the NOAA/NSIDC Climate Data Record of Passive Microwave Sea Ice Concentration (Version 5), and SST from the ESA Sea Surface Temperature Climate Change Initiative (SST_cci) Climate Data Record (Version 3.0).

### Data Processing and Statistical Analysis

Peak areas were normalized by total ion chromatogram (TIC) to estimate relative lipid abundance per unit mass. Data were log-transformed and mean-centered. To avoid division errors, zero signals in both analyte and internal standard channels were imputed as half the lowest detected non-zero value per compound (“half-LOD”).

Class-specific ionization efficiency differences were corrected using the EquiSPLASH internal standard (Avanti Polar Lipids, Alabaster, AL, USA; Supplementary Table S3). Correction factors were calculated as the ratio of the global maximum response to the median internal standard signal for each lipid class. To limit over-correction from outliers, correction factors were capped at ±20% of the compound median. To assess ionization efficiency of wax esters, which did not have an analogous internal standard, two commercial standards (palmitoleyl oleate and arachidyl arachidonate) were purchased (Nu-Chek Prep, Elysian, MN, USA) and made up as quantitative standard mixes. The standards were measured and dissolved in mixtures of methanol and IPA (1:1, v/v) with 0.2 – 0.4mL of DCM added to enable dissolution. The two wax ester standards represented the lower (palmitoleyl oleate; 504.49 Da) and middle (arachidyl arachidonate; 584.55 Da) mass range of the wax esters present in the copepod sample set (418 – 786 Da). Wax ester standards in the higher mass range were not commercially available. Other lipid classes that did not have adequate representative internal standards retained their original values.

Hierarchical clustering was performed on the normalized dataset using the scipy package in Python 3.8. Correlation distance and average linkage were used to cluster both samples and compounds. Clustering methods were selected to emphasize co-regulation patterns in lipid profiles. Principal component analysis (PCA) was used to summarize among-sample variation in fatty acid composition. For each sample, the abundance of each fatty acid was obtained by summing the normalized abundances of all annotated lipid species containing that fatty acid, with fatty acid identities assigned from MS2 fragmentation; lipids lacking fragmentation-level FA information were therefore excluded. PCA was conducted using the PCA function from the sklearn.decomposition module in Python.

## Results

### Zooplankton sampled from three oceanographic regions with varying input from the Atlantic

Distinct water masses occupy each fjord, strongly influenced by the physical oceanography of the surrounding Greenland Sea. Kongsfjord (KB3), the northernmost fjord (Figure 1A), is primarily influenced by North Atlantic waters, characterized by warmer surface temperatures year-round (Figure 2A). In contrast, Storfjorden (SF), located further south, is dominated by Arctic water masses and exhibits consistently colder surface waters (Figure 2D). Billefjord (BAB), situated between KB3 and SF, experiences a combination of these water masses (Figure 1C). Temperature-salinity (T-S) diagrams effectively illustrate these hydrographic distinctions (Figure 3). At the time of sampling, copepods from KB3 were captured within Atlantic Warm water (AWw), those from BAB within Greenland Sea Deep Water (GSDW), while SF copepods traversed multiple water masses, including Fresh Atlantic water (AWf), Atlantic Cold water (AWc), and Eurasian Basin Deep Water (EBDW) [68, 69]. Oxygen and temperature profiles (Figure 1D, E) indicated possible intrusions of warmer, saline waters at approximately 50 m depth in BAB and at 100 m depth in SF, highlighting complex subsurface dynamics.

**Figure 1.**
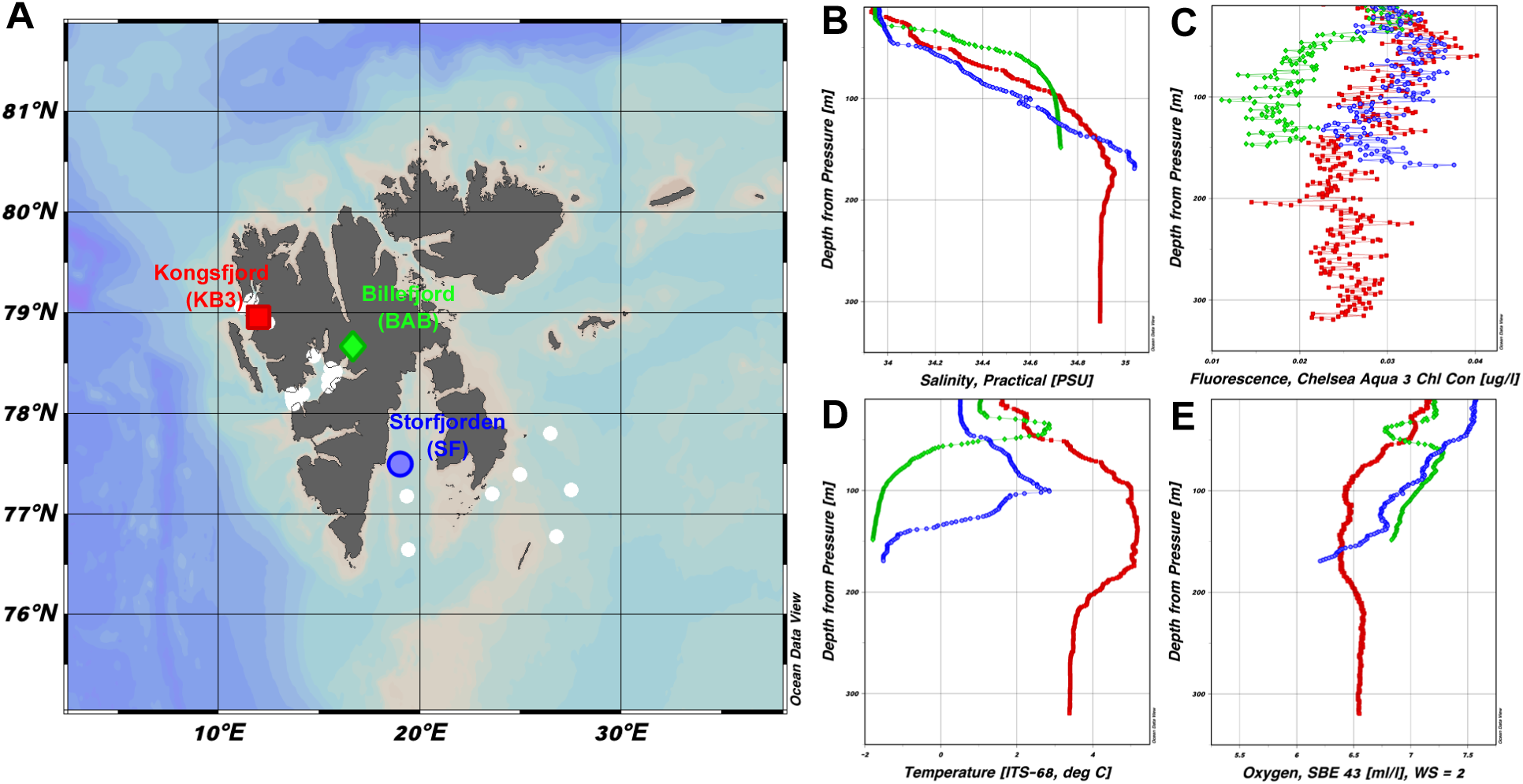
CTD profiles and station locations in Svalbard fjords during the November 2022 cruise. (A) Map of the three focal sampling sites: Kongsfjord (KB3, red square), Billefjord (BAB, green diamond), and Storfjorden (SF, blue circle). White filled circles indicate additional CTD stations conducted during the cruise. (B–E) Depth profiles of (B) salinity, (C) chlorophyll-a fluorescence, (D) temperature, and (E) dissolved oxygen for the three sites, with colors corresponding to panel A (red: KB3, green: BAB, blue: SF).

**Figure 2.**
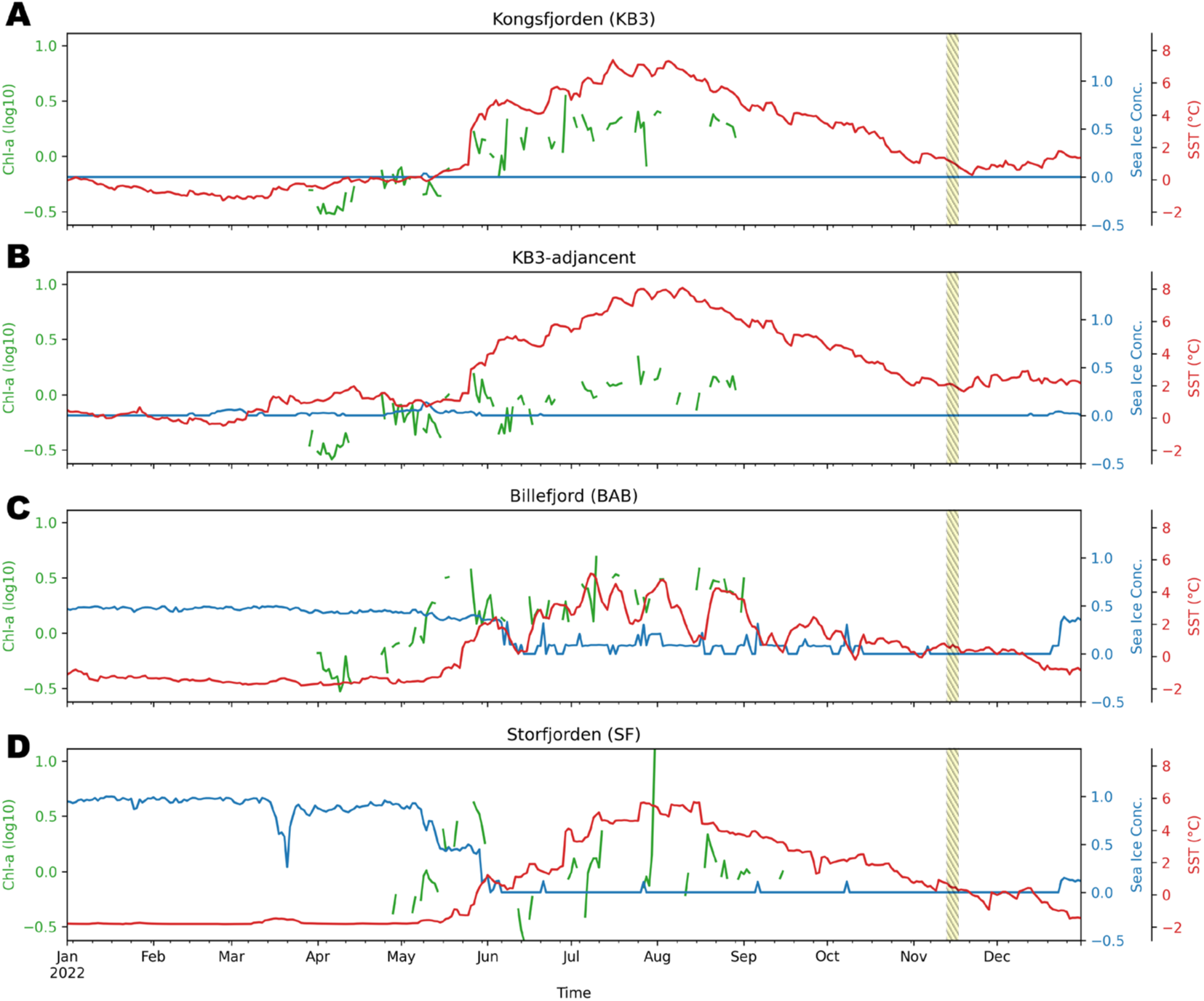
**Temporal variation in chlorophyll-a (log₁₀; green), sea ice concentration (blue), and sea surface temperature (SST; red) at four locations in Svalbard in 2022**: (A) Kongsfjord (KB3), (B) a KB3-adjacent offshore site, (C) Billefjord (BAB), and (D) Storfjorden (SF). All datasets represent daily measurements averaged over a ±0.5° latitude and ±1° longitude spatial window (BAB: 78–79°N, 13–19°E). The KB3-adjacent site was selected to mitigate coastal interference in satellite retrievals at KB3, providing a more representative measure of offshore conditions. The shaded yellow region indicates the field sampling period (13–17 November 2022).

**Figure 3.**
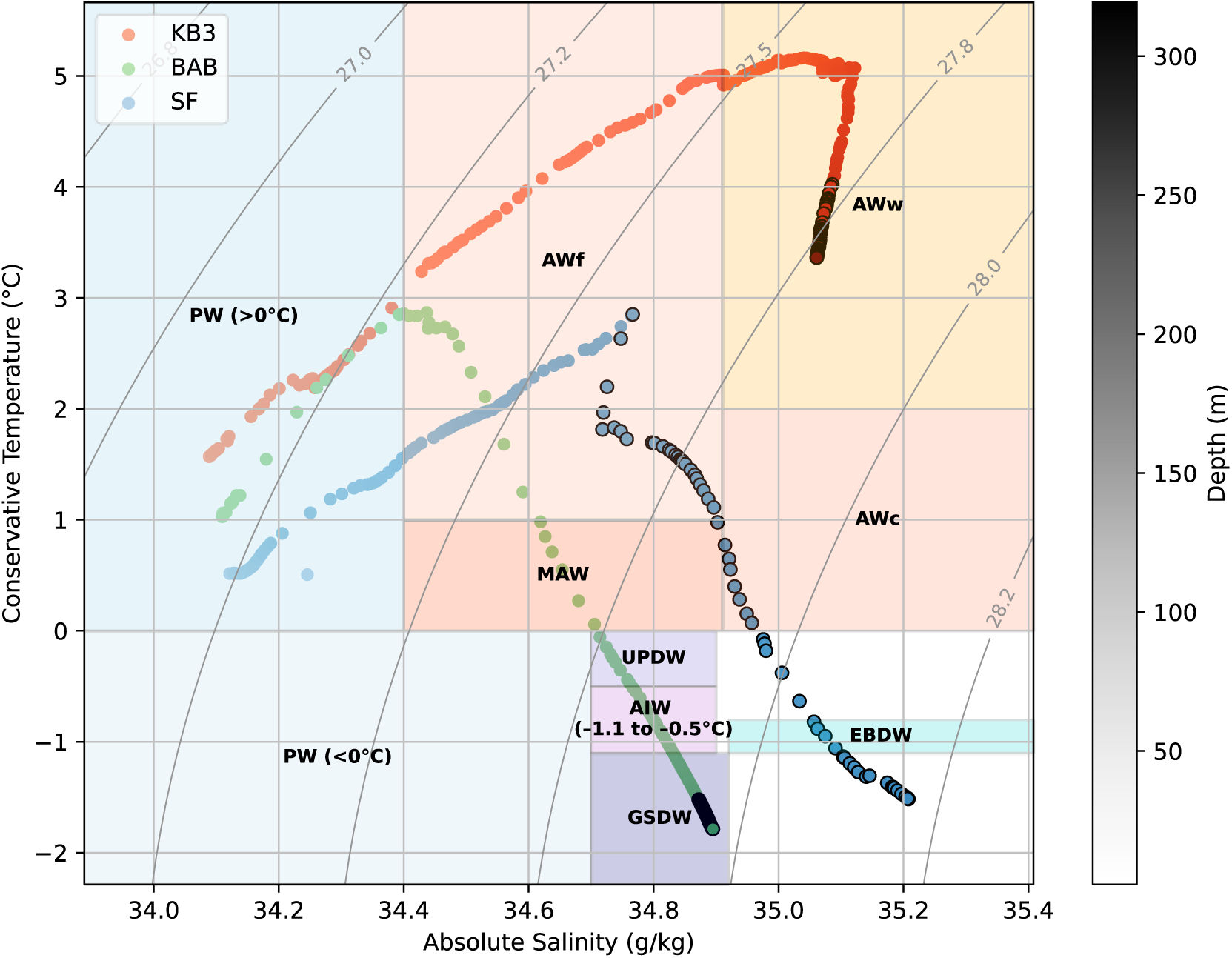
**Temperature–salinity (T–S) diagrams**. Points are shaded by depth, and copepod sampling depths outlined in black. Isopycnals (gray lines) represent potential density anomaly (σ₀, kg m⁻³). Water mass definitions follow Schlichtholz & Houssais (2002), based on the θ–S correlations in Fram Strait (MIZEX 84 data), providing a classification specific to Fram Strait hydrography. An alternative classification using the broader textbook framework of Talley et al. (2011) (source: Aagaard, Swift, & Carmack, 1985) is provided in Supplementary Figure S1.

### Temporal patterns in sea-ice, chlorophyll, and temperature show signs of intensified warming at Kongsfjord relative to other fjords

#### Annual trends for 2022

Sea surface temperatures (SST) and sea ice coverage varied significantly among the fjords throughout the year (Figure 2). KB3 had the warmest SST and remained ice-free (Figure 2A), while SF experienced extensive ice coverage lasting well into May (Figure 2D). Satellite-derived chlorophyll-a (chl-a) data indicated phytoplankton blooms initiating in early April across the region, except in SF, where persistent sea ice delayed bloom onset until May. The simultaneous occurrence of elevated chl-a signals with sea ice melt in BAB and SF suggests potential bloom formation associated with ice-edge dynamics or ice algae presence, though satellite detection of ice algae is challenging [4] (Figure 2C, D). Although SF blooms commenced later and were shorter in duration compared to KB3 and BAB, their intensity was notably higher, likely due to nutrient enrichment associated with cold Arctic inflows. BAB exhibited highly variable hydrographic conditions, reflected by fluctuations in sea ice and temperature (Figure 2C). At the time of sampling (indicated in yellow in Figure 2), all sites were ice-free. Satellite records indicated the cessation of phytoplankton blooms in late August at KB3, early September at BAB, and mid-September at SF.

#### Decadal trends

Satellite observations over the past two decades highlight marked differences in long-term sea ice dynamics among fjords (Figure 4). KB3 has experienced significant reductions in sea ice coverage, indicative of “Atlantification,” the increased influence of warmer Atlantic waters within Arctic environments (Figure 3, 4A, 4B). BAB has also exhibited notable sea ice loss, though less pronounced than KB3 (Figure 4C). Conversely, SF has maintained relatively consistent sea ice coverage, representing conditions characteristic of traditional Arctic marine systems (Figure 4D).

**Figure 4.**
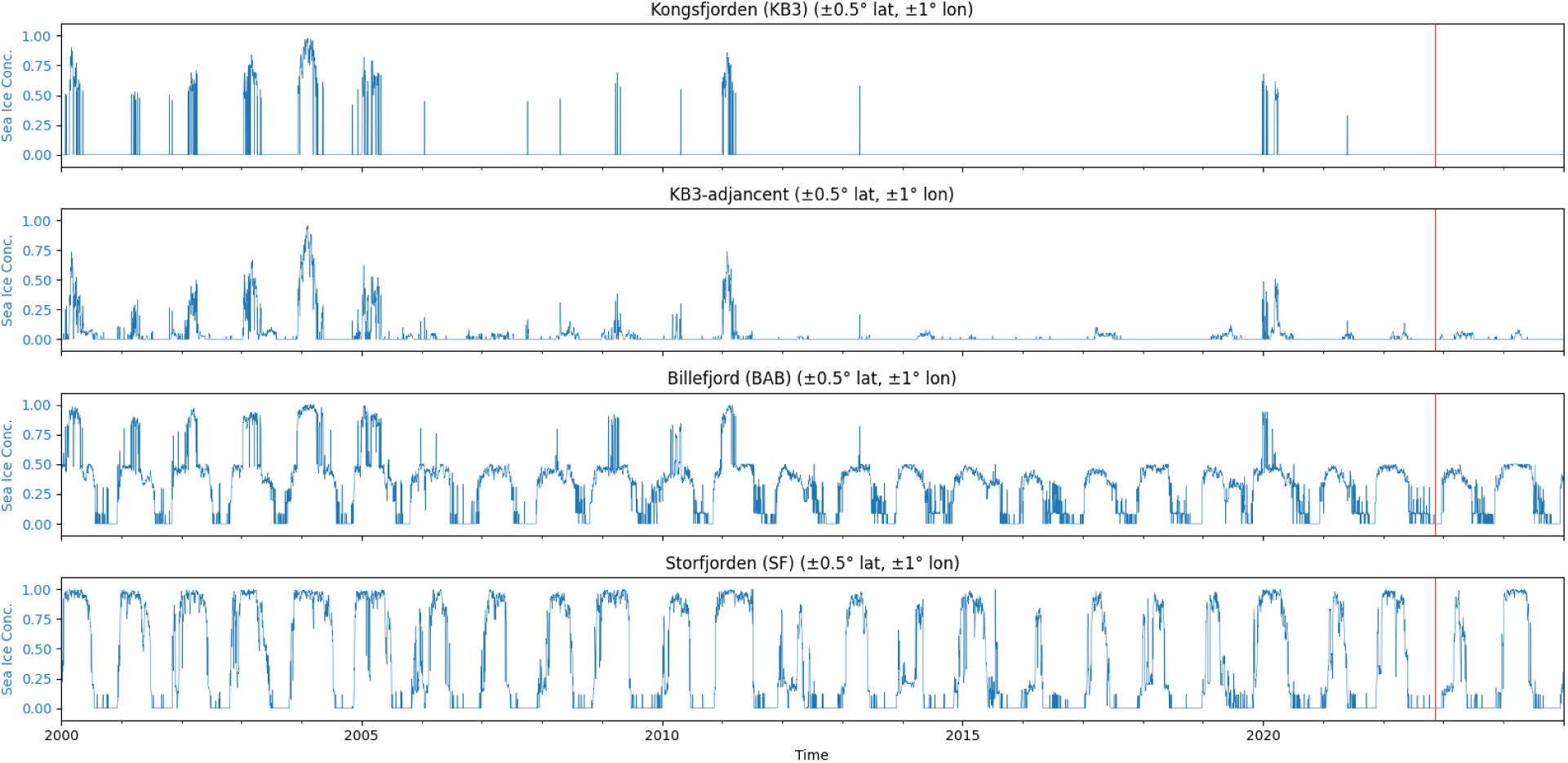
Long-term daily sea ice concentration (2000–present) for three Svalbard fjords: Kongsfjord (KB3), a KB3-adjacent offshore site, Billefjord (BAB), and Storfjorden (SF). Sea ice concentration data are from the NOAA/NSIDC Climate Data Record of Passive Microwave Sea Ice Concentration (Version 5) and averaged over a ±0.5° latitude and ±1° longitude window (BAB: 78–79°N, 13–19°E). The red line indicates the field sampling period (13–17 November 2022).

### Copepodite stage V and individuals collected in Storfjorden had the highest total lipid content

TIC (total ion current; defined here as the summed peak area across all detected lipid features as a proxy for total lipid content) varied significantly among copepods from different fjords (Figure 5), corresponding to the thermal gradients observed. SF copepods, inhabiting the coldest waters, had the largest lipid reserves. The most pronounced size difference was between KB3 copepodite stage 4 (CIV) individuals and SF copepodite stage 5 (CV) individuals, with the latter approximately five times richer in lipid content (Figure 5).

**Figure 5.**
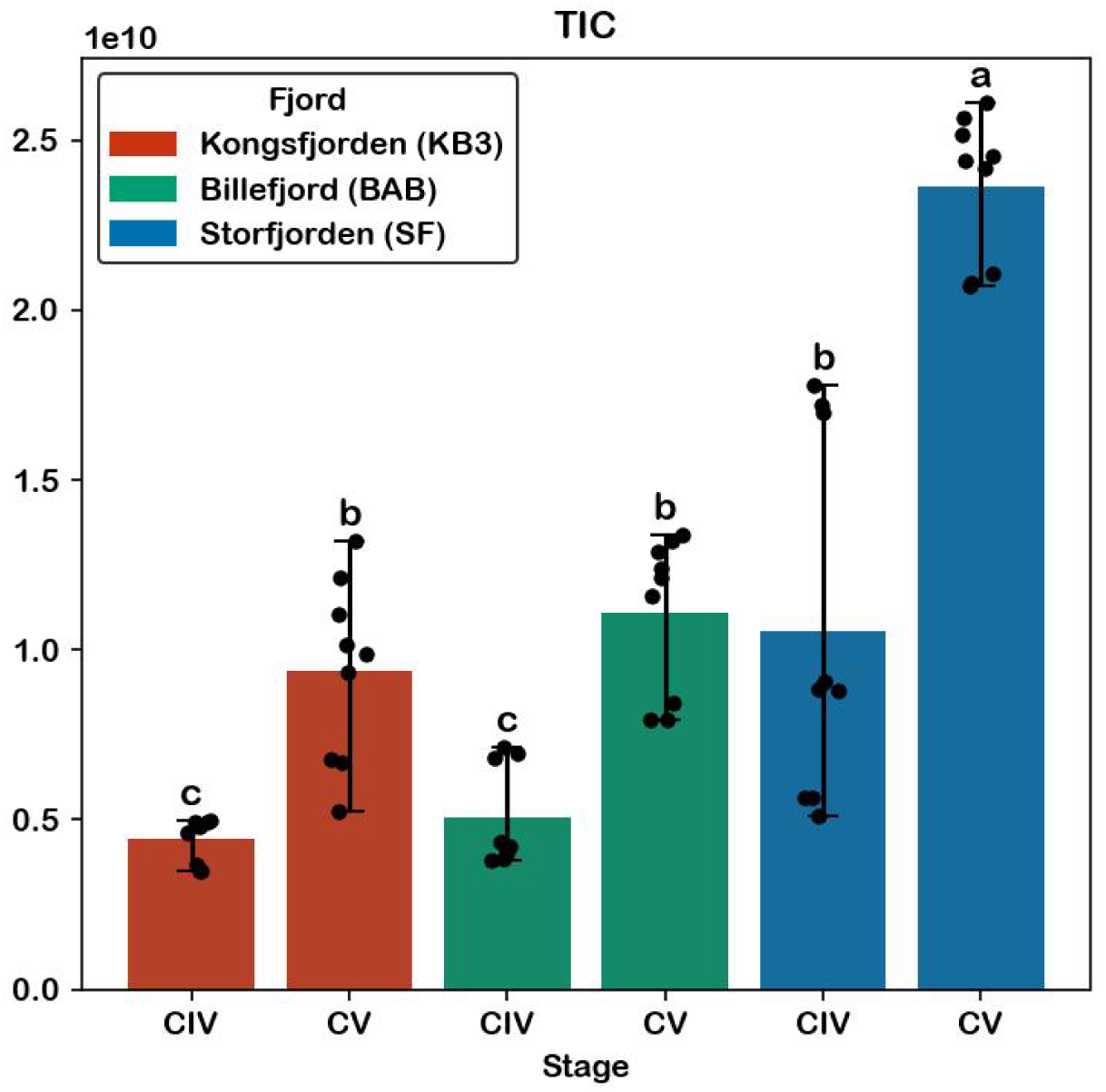
**Total ion current (TIC) across samples**. TIC represents the summed ion signal intensity across all detected compounds in each sample, providing an integrated measure of total extractable lipid abundance. Bars represent the mean TIC for each sample group, with colors indicating fjord identity (red: Kongsfjord (KB3), green: Billefjord (BAB), blue: Storfjorden (SF)). Error bars indicate minimum–maximum ranges, and black points represent individual biological replicates. TIC values have been normalized for recovery and blank-corrected. Group differences were assessed using one-way ANOVA (F = 54.40, p = 1.27 × 10⁻¹⁸) followed fac by Tukey’s honestly significant difference (HSD) post hoc test (α = 0.05). Different letters denote statistically significant differences among groups based on Tukey HSD.

The difference in biomass between fjords was exacerbated when comparing amount of carbon per individual in the form of storage lipids (Figure 6, Table 1). For example, while TIC for SF CV individuals was roughly 5.4-fold larger than that of KB3 CIV, storage lipid carbon content was larger by approximately 19.2-fold.

**Figure 6.**
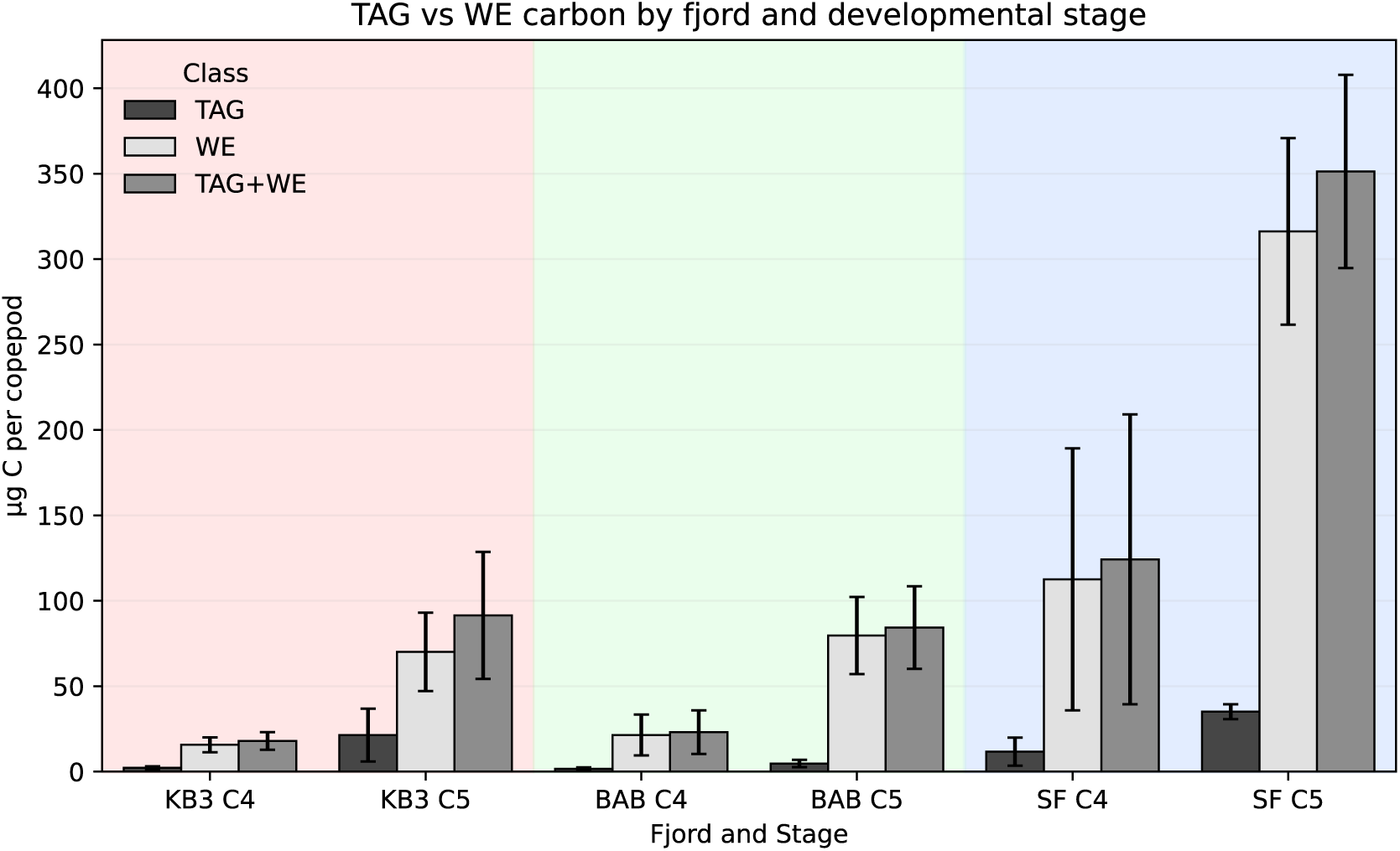
Triacylglycerol (TAG) and wax ester (WE) - derived carbon content per copepod across fjords and developmental stages. For each sample, TAG- and WE-associated carbon (µg C per copepod) were calculated and summed to obtain total neutral lipid carbon (TAG+WE). Bars show mean values for each fjord × stage group, with error bars indicating standard deviation. Shaded background regions delineate fjords (KB3: red, BAB: green, SF: blue), and x-axis labels indicate fjord and copepod stage.

**Table 1.** Comparison of potential contribution of diapausing copepods to biological carbon pump based on wax ester content.

|  | **WE carbon per individual (ugC/ind)** | **Neutral lipid carbon (WE + TAG) per individual** | **Annual contribution of WE to BCP (gC/m<sup>2</sup>/yr)** | **Application to eastern Norwegian Sea (ktonC/yr)** |
| --- | --- | --- | --- | --- |
| **Sinking flux** | - | - | 2 – 8 | 366 – 1464 |
| **Surveys from 2006 – 2008** | 158 | - | 3.2 – 3.7 | 586 – 677 |
| **November 2022 (SF)** | 227.7 ± 139.2 | 251.1 ± 153.0 | 5.3 ± 3.3 | 976 ± 596 |
| **November 2022 (KB3)** | 45.6 ± 36.0 | 57.4 ± 50.5 | 1.1 ± 0.8 | 195 ± 154 |

Values for the November 2022 samples are reported as mean ± standard deviation across three biological replicates (n = 3), each a pooled extract of three individuals; the three analytical replicates per biological replicate were averaged before computing the standard deviation to avoid pseudoreplication. Colors reflect Arctic (blue) and Atlantification (red) scenarios. Values for sinking flux, 2006 – 2008 surveys, and Norwegian Sea surface area are from Jónasdóttir et al., 2015. [15]

Quantification of neutral-lipid carbon per copepod reinforced these fjord-level differences (Figure 6, Table 1). Wax esters dominated the storage pool in every group, accounting for 67–91% of combined TAG + WE carbon. KB3 copepods held the smallest reserves, averaging ∼65 µg C per copepod in CV and ∼12 µg C per copepod in CIV, followed by BAB (∼74 and ∼15 µg C per copepod), while SF carried by far the largest (∼232 and ∼82 µg C per copepod). Within each fjord, CV individuals contained more storage carbon than CIV. The largest difference was between SF CV and KB3 CIV, which differed roughly 19-fold in storage-lipid carbon despite a ∼5.4-fold difference in TIC. KB3 CV showed the lowest wax ester fraction (67%), compared with ≥85% in all other groups.

### Copepod lipidome composition linked to fjord of origin and stage of development, enriched in TAGs and phospholipids, and depleted in WE

Notable differences were observed in lipid profiles among sampled copepods (Figure 7). Wax esters constituted between 36% and 52% of total lipids across samples (Figure 7A). CIV and CV copepodites from KB3 exhibited the lowest average WE content (∼38%), while SF copepods showed the highest average WE content (∼48%).

**Figure 7.**
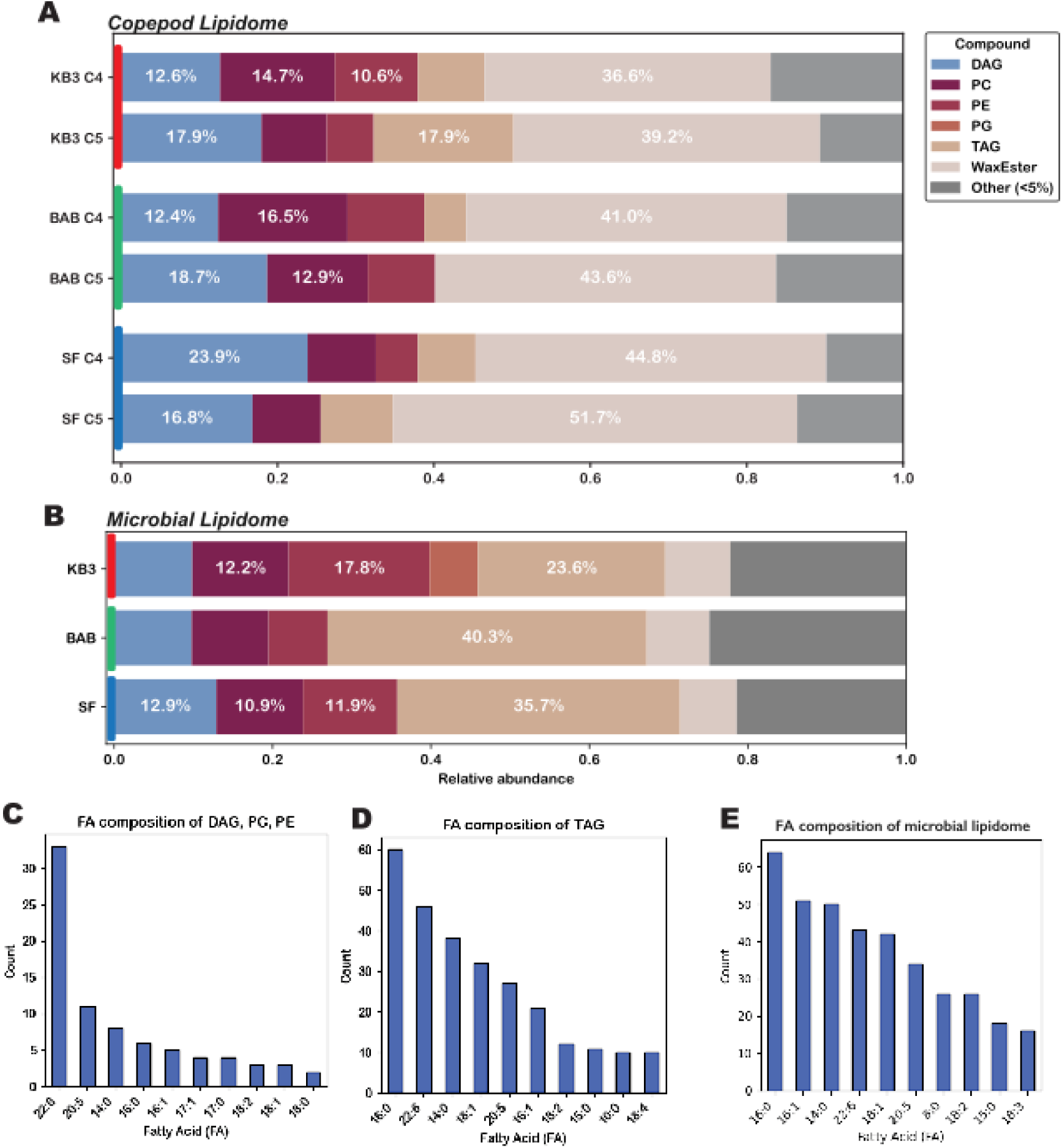
(A-B) Relative abundance of (A) copepod lipid species and (B) surface (10. – **50 m) microbial lipid species.** Bars represent the fractional contribution of each lipid species (colors) to the total lipid pool within each sample group. Only species exceeding a relative abundance threshold (≥3%) in at least one group are shown individually; all others are grouped into “Other” (grey). Values are averaged across replicates and expressed as proportions of the total normalized, blank-corrected lipid signal (TIC). The percentages within each bar indicate the contribution of major species (≥8%). Fjords are ordered from most to least Atlantified: Kongsfjord (KB3), Billefjord (BAB), and Storfjorden (SF). For the surface microbial lipidome, values represent the average composition of the 10m and 50m sample groups. **(C-E) Histogram of the ten most abundant fatty acids composing the (C) major polar lipids and (D) triacylglycerols in the copepod lipidome, and (E) the entire microbial lipidome.**

An additional distinct finding for copepod lipid ecology was the relatively high proportion (>40%) of polar lipids, including phospholipids and diacylglycerols (DAGs) which are not typically reported for overwintering/winter copepods. Compositional fatty acid analysis of these polar lipids (Figure 7C) revealed that many of them were composed of docosahexaenoic acid (DHA), a recognized biomarker for dinoflagellate consumption, suggesting copepods from these fjords might rely more heavily on dinoflagellate-dominated diets than previously reported diatom-dominated diets [70].

Triacylglycerol (TAG) levels exhibited notable spatial variability (Figure 7A). TAG concentrations were high in KB3 CV individuals, accounting for approximately 17.9% of TIC whereas BAB had the lowest TAG levels (below 6%). SF exhibited intermediate, stable TAG levels across developmental stages. This pattern was also reflected in the total fatty acid chain lengths of the lipids weighted by abundance (Supplementary Figure S1). On average, CV individuals had chains approximately 0.8 carbons longer than CIV individuals. Notably, KB3 CV, the copepod group with the most TAGs, showed the longest overall chains, approximately 2.7 carbons longer than their corresponding CIV counterparts.

Methodological considerations should be noted, particularly concerning comparability with previous lipid analyses. Our utilization of UPLC-MS/MS to quantify intact lipids contrasts with traditional lipid analytical approaches employing thin-layer chromatography, fatty acid derivatization methods, and quantification by GC/MS [71]. Although standards have been rigorously tested to minimize ionization bias, methodological differences should be noted while conducting direct comparisons with prior literature.

Hierarchical clustering analysis revealed clear lipidomic patterns related to environmental conditions and copepod developmental stages (Figure 8). A large fraction of lipid compounds (clusters 1 and 2; Figure 8A) were enriched in SF copepods even after normalization for body mass using total ion current. Intriguingly, KB3 CV individuals shared significant lipidomic similarities with SF copepods (namely the dominant cluster 2), despite an apparent contrast in physical environment. This pattern was further supported by hierarchical clustering of the samples (Figure 8B), which grouped KB3 CV individuals exclusively with SF copepods within the same cluster.

**Figure 8.**
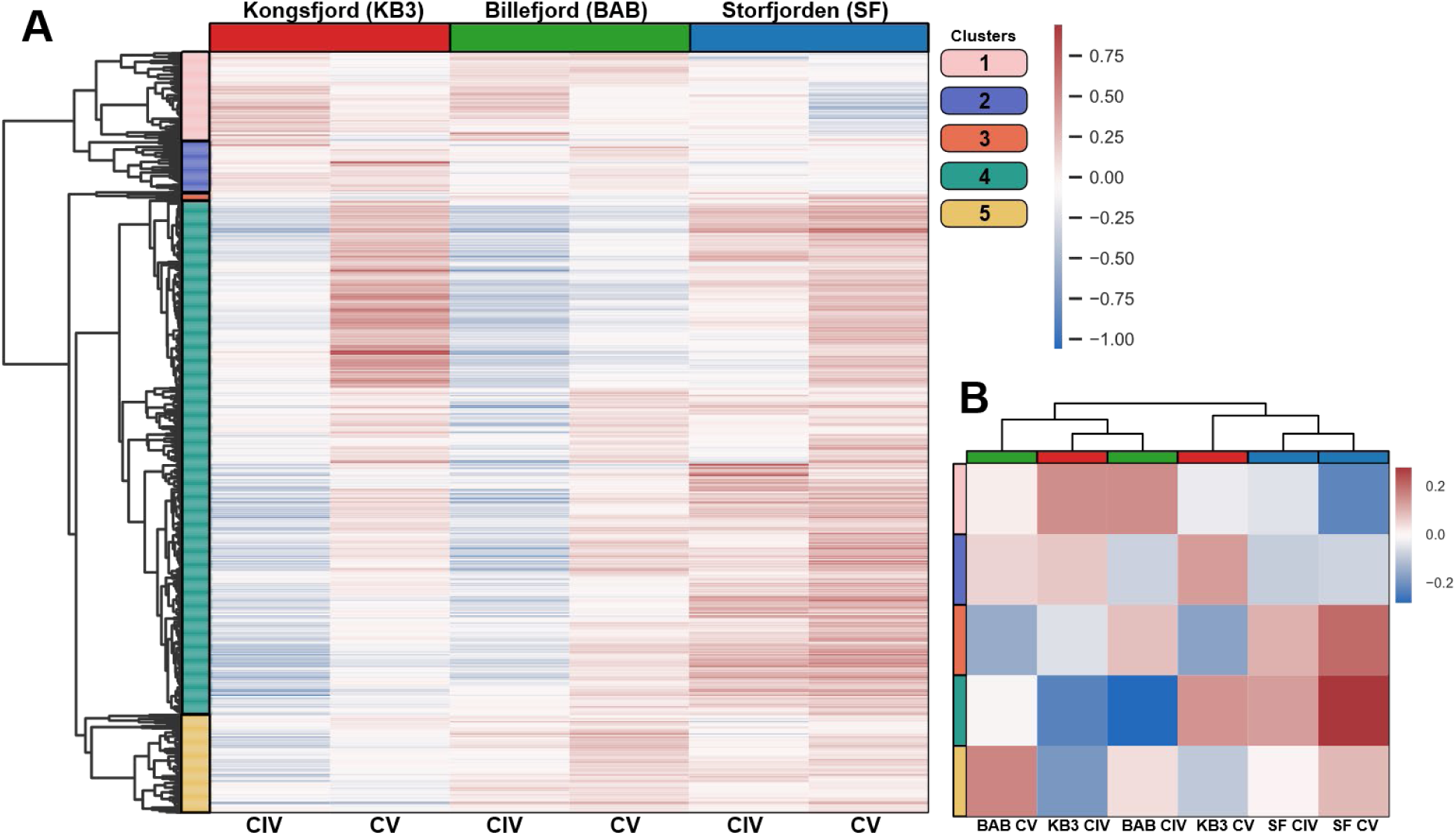
(A) Hierarchical clustering of lipid molecular species detected in copepod samples. Data represent log-transformed, mean-centered normalized peak areas of lipid compounds, averaged across biological replicates within each fjord and life stage group. Fjords are denoted Kongsfjord (KB3), Billefjord (BAB), and Storfjorden (SF). Rows (compounds) are clustered using correlation distance and average linkage into 10 clusters. **(B) Hierarchical clustering of samples using the same data and clustering methods.** Heatmap data represents the average of each cluster-sample pair.

### Lipid analysis of surface microbes hints at link between trophic levels in the winter

Analysis of suspended particulate lipids collected on a 0.7 μm GF/F filter revealed higher compositional variability compared to copepod lipid samples (Figure 7A, B), although the major lipid groups were similar with large contributions from TAGs and phospholipids. WE were less abundant in the microbial lipidome than in the copepods (∼8% of average vs. maximum of 51.7% in SF CV copepodites). Principal component analysis was used to describe the fatty acid composition of the copepod meta-lipidome with principal component 1 capturing 72.9% of the variability in the data and showing 22:6 fatty acid was associated with BAB copepods. Principal component 2 captured only 12.8% of the variability, showing that 16:0 fatty acid was associated with KB3 copepods and 20:5 was associated with SF copepods (Figure 9A). Similar analysis on the microbial lipidome showed 22:6 was associated with the surface microbiome of BAB and SF, whereas 16:n fatty acids and oxidized 20:5 fatty acids were associated with KB3 (Figure 9B).

**Figure 9.**
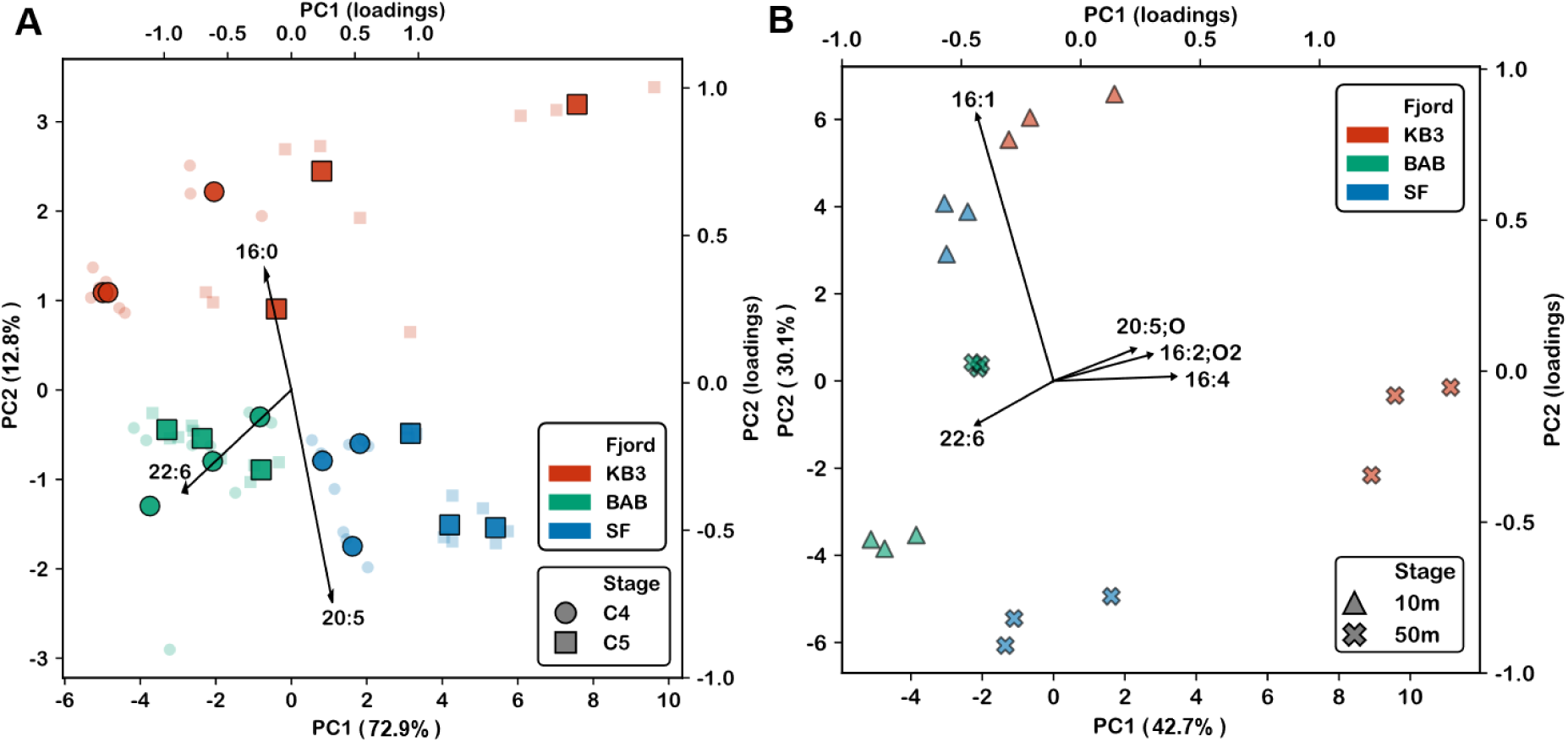
Principal component analysis (PCA) of fatty acid composition of (A) copepods and (B) microbial biomass. Sample points are colored by fjord (red = KB3, green = BAB, blue = SF) and symbol shapes denote copepod developmental stage or depth (circle = CIV, square = CV, triangle = 10m, cross = 50m). Arrows indicate loadings for significant fatty acids (|loading| > 0.3), showing the direction and magnitude of their contribution to the separation of samples along the principal components. In the copepod analysis, analytical replicates (small faint points) are projected post-hoc into the same PCA space for reference. There were no biological replicates for the microbial samples. PC1 and PC2 explained 72.9% and 12.8% of the total variance, and in the microbial, 42.7% and 30.1%, respectively.

## Discussion

### Atlantification led to reduced accumulation of WE in addition to lower overall lipid content

Our findings highlight key biochemical and ecological signatures of climate-driven change in Arctic marine ecosystems, revealing how shifts in ocean temperature, sea ice regimes, water mass structure, and connected food webs are reshaping the phenotypic biology of *Calanus* copepods. Specifically, *C. glacialis* copepodites in the CIV and CV stage had unusual lipid compositions for diapausing species (Figure 7A) and individuals from Kongsfjord, which was mostly Atlantic Warm water at depth, had significantly lower lipid content (Figure 5, 6), and a skewed lipid composition compared to the most ice-covered (Figure 7A), Arctic-influenced site, Storfjorden which had the highest lipid content and more typical lipid composition for wintertime (i.e. higher relative abundance of WE; Figure 5-7). Our discussion below underscores the potential for Atlantification to alter lipid storage, feeding strategies, and carbon transport, with implications for the broader Arctic food web and biogeochemical cycles.

Wax esters are critical for energy storage and buoyancy regulation during diapause in *Calanus* species [22, 48]. The WE proportions observed in this study (36 – 52%; Figure 7A) are substantially lower than the ranges typically reported for *Calanus glacialis* (70–90%) in previous studies [18, 19, 21, 22, 62, 70]. The low WE content observed at KB3 is comparable to lipid levels reported for actively feeding summer copepods rather than diapausing individuals [72]. In contrast, the relatively higher WE content observed in SF copepods is more consistent with expectations for the colder environmental conditions observed there due to the larger Arctic/EBDW influence and duration of sea ice coverage (Figures 2, 3, 7A). Reports of wax ester content in diapausing *C. glacialis* are limited [73, 75], with most existing studies conducted during spring, summer, or fall when individuals are actively feeding. By comparison, data for the sympatric diapausing *C. finmarchicus* are more abundant, with reported WE proportions ranging from 80–99% at depths and latitudes comparable to those of the present study [57].

Therefore, we consider KB3 copepods the “Atlantification” phenotype, SF the “Arctic” phenotype, and BAB as an intermediate. If Atlantification becomes the dominant process in the Arctic, we would anticipate the promotion of KB3-phenotype *C. glacialis* across the entire Arctic shelf. Before we discuss the quantitative implications for the seasonal lipid pump, we compare our detailed lipidomic results with other studies investigating the effects of Atlantification on lipid storage and size, and we explore two other factors that likely contribute to the “Atlantification” and “Arctic” meta-lipidomics signals: (1) winter feeding and carnivory, and (2) population origin and movement.

### Effects of Atlantification on lipid storage and individual size

Warmer water temperatures influence copepod development, evident in lower lipid reserves and overall size, as reflected in the total ion current and wax ester content among our samples (Figures 5, 6, 7). Although elevated metabolic rates in warmer environments may partly explain reduced lipid reserves [59], the fact that TAGs are typically used up faster than WE during starvation or diapause [70] makes their high abundance in the “Atlantified” KB3 notable (17.9% of the lipidome; Figure 7). This is also in spite of the depth dependence of lipid concentrations observed in Antarctica where Pond (2012) reported an average increase of 100 µg of total lipid per individual with every 100 m depth below 200m [31]. Yet KB3 copepods sampled from 200–330 m still exhibited lower WE levels compared to SF and BAB, which were sampled above 200 m (Figure 6; *C. glacialis* diapause depth is 200 – 500 m). Assuming the copepod specimens were in true diapause, the disproportionately high triacylglycerol levels contrasted with the lower WE storage in KB3 would indicate potential differences arising from diet during the spring lipid accumulation phase.

Our findings corroborate predictions from Aarflot et al. (2023) and a 13-year time series by Møller and Nielsen (2020), linking higher temperatures and altered food availability to reduced copepod size and lipid content [9, 23]. Increased metabolic rates may further accelerate lipid depletion through respiration, potentially reducing the lipid flux to higher trophic levels and altering diapause duration [59]. This is also consistent with hypotheses posed by Pierson et al. (2013), who suggested that climate change would likely have differing effects on a species across its geographical range, although they did not have the empirical data needed to evaluate the potential impact that phenotypic variability has on dormancy duration.

An additional consequence of Atlantification is the potential for increased size niche overlap between *Calanus finmarchicus* and *C. glacialis*. Weydmann et al. (2022) previously documented significant niche overlap based on population dynamics (population size and stage ratios) correlated with environmental metadata [73], rather than lipid-based indicators as we have done here. However, it is still an unresolved question whether *C. finmarchicus* can sustain self-recruiting populations in Svalbard fjords and the Barents Sea over multiple years. In particular, Arctic winter conditions may act as a demographic “filter” by constraining overwintering survival and completion of their life cycle. If winter filtering is weak, recurrent co-occurrence could promote a community-level shift toward boreal, *C. finmarchicus*-like dominance, consistent with the borealization of Arctic zooplankton documented elsewhere [9, 11]. Whether such a shift would also involve genetic exchange remains unresolved: natural hybridization between the two species has been reported [74] but was not supported by subsequent SNP-based analyses, which found clear species differentiation and no admixture [75]. Conversely, if winter filtering remains strong, recurrent overlap may not translate into long-term replacement. We reiterate that we used classic taxonomic distinguishers for *C. glacialis* and *C. finmarchicus* and CIV and CV, but we do not have the DNA data to ask whether hybridization is at play here. We explore explanations more well backed by existing literature below. However, it is important to note that, typically, *C. glacialis* is visually confused for *C. finmarchicu*s and not the other way around [63].

### Winter Feeding and Carnivory

Another factor that may help explain the distinct KB3 lipidome is winter feeding activity. As mentioned before, diapause is not mandatory, and in the past two decades, there have been several reports of winter activity, diel vertical migration (DVM) and feeding in *Calanus* copepods [26, 70, 76–80]. These studies, using 18S gut content analyses, acoustic Doppler current profiler (ADCP) moorings, and measurements of respiration and swimming activity, concluded that *C. glacialis* individuals can not only remain active but also exhibit flexible feeding strategies, switching to carnivory in winter in addition to bypassing diapause under certain conditions. Krumhansl et al. (2018) suggested such copepods may have emerged from diapause earlier than expected or been advected upward by upwelling and mixing processes [29].

Low lipid stores, especially of wax esters, as we saw in KB3 and BAB, are more typical of non-dormant copepods [72] and have also been singled out as one of the main criteria for active feeding behavior (hunting and carnivory) in these organisms [80]. It has previously been hypothesized that copepods must reach a quota of WE to enter diapause, referred to as the Lipid Accumulation Window; the quota for *C. finmarchicus* is estimated to be roughly 70 μg C as WE in order to enter diapause at stage CV [22]. Although it has not been systematically tested in the field, our low WE percentages would suggest the population may not have met the lipid quota, especially considering *C. glacialis* is larger and more lipid rich compared to *C. finmarchicus* (Figure 6 KB3-CV average 53 μg). In addition, the copepod lipidomes exhibited elevated levels of phospholipids (PC, PE) and glycerolipids (DAG) rich in docosahexaenoic acid (DHA; 22:6), a known dinoflagellate dietary biomarker in copepods [19, 21, 81] (Figure 7C). Kohlbach et al. (2021) similarly linked DHA, and another lipid proxy (the 18:1(n:9)/18:1(n-7) ratio) to winter carnivory in the Barents Sea [70]. While these lipid biomarkers are not unambiguous, we will explore the biochemical and geochemical implications of these potential dietary sources below.

Biochemically, preferential catabolism of dietary long-chain polyunsaturated fatty acids (LC-PUFAs) such as EPA and DHA, along with prioritization of TAG-fueled metabolism during prolonged starvation, typically results in depleted levels of these compounds in diapausing copepods [31, 70]. However, KB3 copepods retained substantial TAG and LC-PUFA content, suggesting recent or ongoing feeding (Figure 7). Future research involving controlled dietary experiments could further elucidate dietary fatty acid turnover and lipid metabolism dynamics during diapause.

Principal component analysis (PCA) of copepodite fatty acid composition showed strong correlations between DHA and BAB samples, palmitic acid (16:0) and KB3 samples, and between eicosapentaenoic acid (EPA; 20:5) and SF samples, implying distinct dietary sources of these lipids [19, 21, 81] (Figure 9). While DHA serves as a dietary biomarker for dinoflagellates and EPA represents a diatom-dominated diet, palmitic acid is ubiquitous across most organisms.

Assuming these lipids accurately reflect dietary sources, the winter lipidome of SF copepods likely resulted from trophic transfer of diatom biomass over the year. In contrast, BAB copepods appear to have fed primarily on dinoflagellates, which may be present across multiple seasons due to their mixotrophic capabilities and were the dominant phytoplankton subclass in September 2022 [82]. Our PCA of microbial biomass indicated that the surface waters of Billefjord (BAB) were highly correlated with DHA, corroborating the presence of winter dinoflagellates (Figure 9). BAB CV copepods wholly lack TAGs (Figure 7A) and the CIV have the next lowest concentration (Figure 6, 7A), combined with the low WE accumulation (Figure 6) and disconnect between high TAG content in surface microbiomes from the same fjord (Figure 7B), we speculate that these copepods may have been exhibiting stress and burning through their TAG reserves. The association of palmitic acid with KB3 copepods suggests a more generalized feeding strategy (Figure 9), potentially reflecting opportunistic grazing during the resource-limited early winter. Previous reports from Billefjord observed gut contents of active copepods in winter which included copepod nauplii and eggs (cannibalism), early-stage chaetognaths, fungi, and dinoflagellates, particularly those observed in fall and early winter [78].

Although a direct link between winter feeding and warming has not been proven, our observations suggest that this behavior likely contributes to the lipid composition seen in KB3 copepods (Figure 5, 6, 7). Warmer sea temperatures can speed up the use of wax ester reserves in diapausing individuals, potentially forcing them to return to surface waters earlier or more often to feed, as shown in studies on temperature and diapause duration. Based on these lipid profiles, we suggest that some populations, especially those in KB3 and BAB, are not fully diapausing but are moving up to the surface to feed on, in the case of BAB, what seems to be a dinoflagellate-rich food source and in the case of KB3, on opportunistically available particulate resources.

### Population Origin and Movement

The unique lipid composition observed in KB3 copepodite stage IV individuals compared to stage V could also suggest non-endemic origins; that these copepods may be ecological “expats” that originated elsewhere before arriving in the sampling area. Although the specificity to stage is unusual, it could be explained by the fact that CIV and CV do not descend at the same time and later descent typically resulting in diapause at a later stage [23]. Aarflot et al. (2023) noted uncertainties about *C. glacialis* population sources in the Barents Sea, while Scott et al. (2000) and Basedow et al. (2018) also suggested advective transport of copepods among fjords [23, 62, 83].

These observations, combined with the uncertainty around molting during winter, highlight key knowledge gaps in developmental timing, life stage transitions, and the possible influence of non-endemic copepod populations on local lipid signatures. These points underscore the need to consider both physical transport and life-history variability when interpreting lipid profiles across fjords. Regardless of the mechanism of the lower lipid reserve phenotype in the most “Atlantified” waters, the predicted consequences are less robust zooplankton populations with lower recruitment and lower metabolic reserve to battle environmental stress.

### Quantitative Assessment of Atlantification Impact

To quantify potential consequences of widespread Atlantification, we conducted a simple back-of-the-envelope calculation that illustrated potential reductions in the magnitude of the seasonal lipid pump (Table 1). Briefly, we quantified WE-derived carbon per CV individual and compared it proportionally to values reported by Jónasdóttir et al. (2015) to estimate how each phenotype would contribute to the seasonal lipid pump across the Eastern Norwegian Sea (see Methods for details). The magnitude of the North Atlantic biological carbon pump (BCP) has been estimated at 2–8 g C m⁻² yr⁻¹. Jónasdóttir et al. (2015) estimated that diapausing copepods contribute an additional ∼3.7 g C m⁻² yr⁻¹, roughly comparable to the BCP itself. In our calculations (Table 1), “Arctic” phenotypes would contribute ∼5.3g C m⁻² yr⁻¹, which remains within the range considered comparable to the BCP and therefore represents a substantial additional flux. By contrast, a predominance of “Atlantified” phenotypes would reduce this contribution to ∼1.1 g C m⁻² yr⁻¹, a 70.2% decline in SLP strength relative to the 2006–2008 survey used in Jónasdóttir et al. (2015). Taken together, these results indicate that a shift toward KB3-type “Atlantified” *C. glacialis* phenotypes, as suggested by multiple observations and models [15, 84–86], would drastically reduce biological carbon sequestration in the Arctic.

Despite growing recognition of the Arctic’s rapid warming, there remains a striking lack of wintertime measurements of copepod physiology, particularly in regions undergoing Atlantification. Previous work, such as Tarrant et al. (2016) in the warmer Alaskan Sea, reported unexpectedly high concentrations of polar lipids (PL), which were initially dismissed as contamination [43]. Our findings suggest an alternative explanation: these PL and TAG values may reflect true biochemical shifts under warming conditions, driven by declines in wax ester storage, inability to transition into diapause, and winter feeding as hypothesized for KB3. This reinterpretation underscores how warming-induced changes in copepod lipid profiles may have been previously overlooked or misattributed.

## Conclusion

Although our dataset represents only a snapshot in space and time, it reveals meaningful patterns in copepod lipid composition that align with broader physical and ecological gradients across fjords. Importantly, these patterns, ranging from altered lipid size and storage lipid content to signs of winter feeding and potential non-endemic origins, demonstrate the plasticity of *C. glacialis* and its capacity to persist in a changing Arctic. The emergence of distinct phenotypes, such as the “Atlantified” KB3 type, suggests that while this species may remain prevalent in future Arctic systems, it may do so with altered roles in food webs and biogeochemical processes.

To better understand these transitions, there is a need for long-term, winter-inclusive monitoring of *Calanus* populations, such as the time series analyses described by Weydmann et al. (2022), paired with integrative approaches [73]. For instance, experimental work similar to that of Tarrant et al. [42], comparing diapausing and non-diapausing copepods, could provide critical insight. Such studies, especially if combined with transcriptomics, would help identify the molecular mechanisms underpinning diapause variability and environmental responsiveness. Ultimately, characterizing these shifting copepod strategies is essential for refining our predictions of Arctic carbon flux and food web dynamics under continued Atlantification

## Acknowledgments

We would like to thank the captain and crew of the Go SARs for enabling sampling collection and the Nansen Legacy project for continuing this critical work. At UC Berkeley, we would like to thank the Peder Sather Center for Advanced Study and the Society of Hellman Fellows for funding this work.

## Acronyms (alphabetical)

AIW: Arctic Intermediate Water
ASW: Arctic Surface Water
AW: Atlantic Water
AWc: Atlantic Cold Water
AWf: Atlantic Fresh Water
Aww: Atlantic Warm Water
DW: Deep Water
EBDW: Eurasian Basin Deep Water
GSDW: Greenland Sea Deep Water
MAW: Modified Atlantic Water
PIW: Polar Intermediate Water
PW: Polar Water
UPDW: Upper Polar Deep Water.

## Notes

### Competing Interest Statement

The authors have declared no competing interest.

