## Supplementary Figure S1 for "Biochemical Indicators of Atlantification and Diapause Strategy in Arctic Copepods Point to a Decrease in Copepod-mediated Carbon Sequestration"

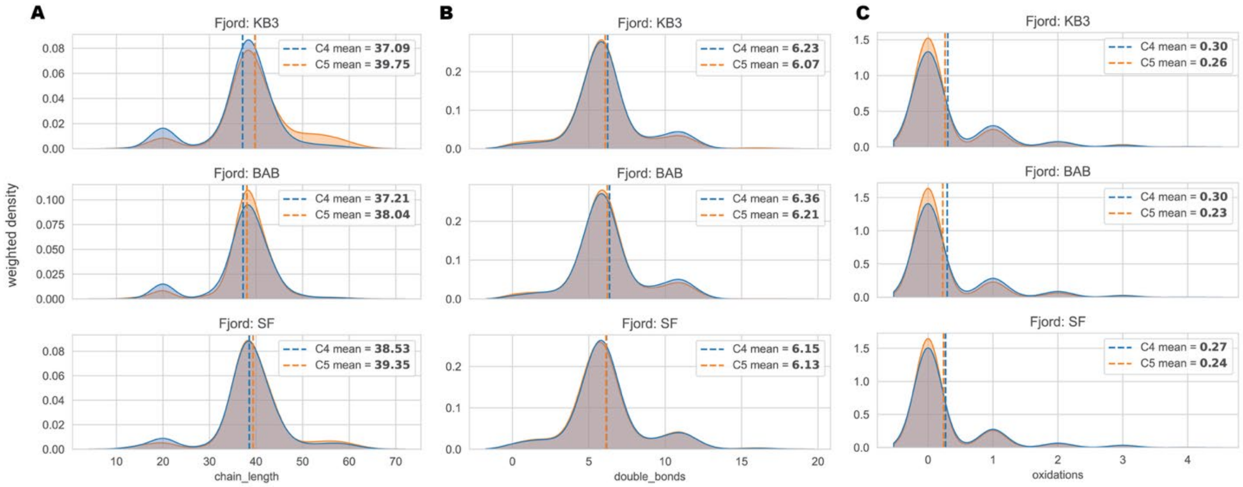


**Supplementary Figure S1**. **Distribution of lipid structural features in copepod lipidomes.** Kernel density estimates are shown for copepod developmental stages CIV and CV, weighted by compound abundance. Panels represent (A) carbon chain length, (B) number of double bonds, and (C) number of oxidations. Dashed vertical lines indicate the abundance-weighted mean for each stage within each fjord.
