## Supplementary Table S1 for "Biochemical Indicators of Atlantification and Diapause Strategy in Arctic Copepods Point to a Decrease in Copepod-mediated Carbon Sequestration"

**Supplementary Table S1. Sampling by site and developmental stage**

| | Sampling depth | CIV | CV |
|---|---|---|---|
| Billefjord | 178 – 100m | 9 | 10 |
| Kongsfjord | 167 – 100m | 11 | 12 |
| Storfjorden | 322 – 200m | 12 | 13 |
