## Supplementary Table S2 for "Biochemical Indicators of Atlantification and Diapause Strategy in Arctic Copepods Point to a Decrease in Copepod-mediated Carbon Sequestration"

**Supplementary Table S2. Data Reduction Table for analysis of data via XCMS-CAMERA-LOBSTAHS-manual verification pipeline**

| Data reduction step | Number of features |
|---|---|
| XCMS-CAMERA Feature detection | 6720 |
| LOBSTAHS annotation | 945 |
| MS-DIAL annotation | 5215 |
| Unique compounds<br>(Manual verification + WE database) | 749 |
