## Supplementary Table S3 for "Biochemical Indicators of Atlantification and Diapause Strategy in Arctic Copepods Point to a Decrease in Copepod-mediated Carbon Sequestration"

**Supplementary Table S3. Ionization-efficiency normalization factors by lipid class.**

Lipid signal intensities were corrected for class-dependent differences in electrospray ionization efficiency using the EquiSPLASH internal standard mixture. For each standard, the best-ionizing adduct was selected, and a normalization factor was computed per sample as the ratio of the reference standard signal to that standard's signal. PC, the highest-responding standard, served as the reference and was assigned a factor of 1.0. The median factor across all 45 samples (shown) was applied uniformly to every compound in the corresponding class. Higher factors indicate poorer ionization efficiency and therefore a larger upward correction. Zero values were imputed at half the lowest detected signal prior to factor calculation. Compounds not listed do not have analogous compounds within the standard mix and hence were not corrected.

| Internal standard | Measured lipid classes normalized to it | Median factor |
|---|---|---|
| PC | PC, EtherPC | 1.0 |
| SM | SM, ASM | 1.1 |
| PE | PE, EtherPE | 1.3 |
| Ceramide | Cer_NS, Cer_HS | 1.5 |
| TAG | TAG, OxTG, VAE | 4.9 |
| PG | PG | 5.0 |
| LPC | LPC | 5.2 |
| — [a] | WaxEster | 15.5 |
| LPE | LPE, EtherLPE, LPA | 12.3 |
| PI | PI | 23.3 |
| DAG | DAG, DG, EtherDG | 48.4 |
| MAG | MAG, FFA | 285.1 |

**Notes**

a Wax esters have no EquiSPLASH analog; their factor (15.5) was derived separately by regression against a custom wax ester reference standard.

b Phosphatidylserine (PS) and cholesterol/coprostanol esters lacked a reliably detected internal standard and were not ionization-corrected.
